# Analysis of structural network topology alterations in autism and its relationship with autism severity

**DOI:** 10.1101/2022.12.20.521280

**Authors:** Abhishek S. Jeyapratap, Drew Parker, Benjamin E. Yerys, Ragini Verma, Yusuf Osmanlıoğlu

## Abstract

Autism Spectrum Disorder (ASD) is a public health concern ranging along a continuum of severity. The neurobiological motives behind ASD have been widely explored with reports about aberrant brain anatomy and functional connectivity. However, research on the underlying structural connectivity alterations is limited. We propose the application of a novel connectomic measure called Network Normality Score (NNS) to identify brain abnormalities and quantity topological dissimilarities in individuals with ASD. We show that the network topology of structural connectivity is altered in ASD brains relative to healthy controls at the global and system levels. We demonstrate that structural connectivity differences are more pronounced in certain subnetworks. Finally, we quantify the association between network similarity and behavioral autism severity to show the efficacy of NNS as a neuroimaging measure.

## 1. INTRODUCTION

Autism Spectrum Disorder (ASD), or Autism, is a public health concern that comprises a set of lifelong neurodevelopmental disorders defined by certain common characteristics such as social and communication issues along with repetitive and restrictive behavioral patterns [1]. Due to the complex etiology and neurobiology of ASD, there is substantial heterogeneity among individuals ranging along a continuum of severity [2].

Diagnosis of Autism is primarily done based on observed behavioral factors [1]. However, due to challenges with distinguishing characteristic behavioral Autism symptoms from typical development and other delays and conditions, ASD can be diagnosed accurately only after 18–24 months of age [3]. The need for earlier diagnosis makes neuroimaging an important candidate to provide reliable markers for the disorder. The neurobiological underpinnings of Autism have been widely explored through neuroimaging tools such as magnetic resonance imaging (MRI), where anatomical differences mainly associated with enlarged brain is reported [4], including increased white matter and cortical surface area as well as abnormal growth patterns in gray matter [5].

Connectomics, analysis of the brain as a network of interconnected regions, has recently become indispensable for the characterization of structural and functional connectivity [6] in healthy brains as well as in brain disorders like ASD [7]. Functional connectivity derived from resting-state fMRI data has demonstrated aberrant connectivity in the ASD functional connectome where long-distance cortical under-connectivity [8], as well short-distance over-connectivity [9] is widely reported. There is also growing evidence for atypical structural connectivity in Autism as measured by diffusion MRI (dMRI). Using scalar measures such as fractional anisotropy (FA) that indirectly quantifies connectivity, reduced FA in frontal and temporal regions [10] can be indicative of reduced structural connectivity in Autism. Although considered a ‘developmental disconnection syndrome’ [11], network-level analysis of structural connectivity in Autism that is derived from dMRI and tractography is very limited.

In this study, we propose the application of a connectomic measure called Network Normality Score (NNS) [12] over Autism to quantify topological differences in the structural connectivity of individuals with ASD (ASDs) relative to a sample of healthy controls (HCs). Using NNS, we first analyze the global differences in brain connectivity of ASDs relative to HCs at the whole connectome level. We then investigate the topological differences at the level of functional systems over frontoparietal and default mode networks. Finally, using an unsupervised clustering approach, we examine the relationship between NNS and Autism severity to determine the association between the structural network wiring and behavioral patterns of autism.

## 2. METHODS

### 2.1. Dataset

In our analysis, we considered a cohort of 150 healthy controls (115 males) and 163 individuals with Autism (133 males) in age range [4,20] (mean=12.1). Individuals with Autism were assessed for the severity of their condition by the standard Autism Diagnostic Observation Schedule (ADOS) [13]. Diffusion MR data was acquired for each subject on a Siemens 3T Verio whole-body scanner with a 32-channel array head coil (single-shot, spin-echo sequence, TR/TE = 11000/76ms, b = 1000s/mm2, 30 directions, 1 [b = 0] volume, flip angle = 180^*◦*^, resolution = 2 × 2 × 2mm). High-resolution T1-weighted anatomic images were also obtained using a 3D MPRAGE imaging sequence with TR = 1900ms, TI = 900ms, TE = 2.54 ms, flip angle = 9^*◦*^, resolution = 0.82 × 0.82 × 0.9mm. Probabilistic tractography was performed over structural MR data and connectomes were generated as a 200 × 200 adjacency matrix of weighted connectivity values by using Schaefer atlas, where each element represents the number of streamlines between regions. The reader is referred to [14] for the details of the data processing pipeline.

In our analysis, we focused on default mode (DMN) and frontoparietal networks (FPN), as these systems were shown in fMRI studies to have altered patterns of connectivity in Autism [4, 9].

### 2.2. Network normality score

To capture the differences in structural network topology and quantify connectomic similarity between Autistic and healthy individuals, we considered a graph matching based approach called Network Normality Score (NNS) presented in [12], which accounts for changes in the global network topology instead of focusing on local connectivity disruptions. NNS is calculated as the average graph similarity of an individual relative to a reference sample. Using healthy controls as a reference, we first calculated NNS among them (denoted NNS_H_). Adopting the measure to Autism, we then calculated NNS of ASDs relative to the healthy control sample (denoted NNS_AH_), which provides a normative connectomic similarity measure.

### 2.3. Statistical Analysis

To evaluate the efficacy of NNS in capturing alterations in the structural wiring of ASDs relative to HCs, we calculated group differences using Welch’s t-test between NNS_H_ and NNS_AH_ at both connectome and system levels. *p*-values are FDR corrected for multiple comparison wherever applicable and effect sizes are calculated using Cohen’s *d*. Since Welch’s t-test is parametric, subjects with outlier scores were discarded from analysis.

Taking an unsupervised learning approach with agglomerative hierarchical clustering, we defined two sub-groups based on each subject’s neuroimaging score as measured by NNS at the connectome and system levels. The number of sub-groups was chosen by inspecting the dendrogram of hierarchical clustering. Using these clustered sub-groups, we assessed the relationship between the similarity of structural network topology of ASD subjects and their Autism severity as quantified by ADOS score with the following linear model (LM) that controls for age:

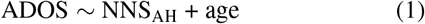

This provides a basis of evaluation for quantifying the effectiveness of NNS as a connectomic measure of Autism since ADOS clinically measures autism severity based on behavioral observations.

## 3. RESULTS

Evaluating NNS of healthy controls and Autistic individuals at the connectome level, we observed a significant group difference between the NNS_H_ and NNS_AH_ distributions (Cohen’s *d* = 0.29, *p* = 0.01). When considered without outliers, significance of group difference would diminish (*d* = 0.20, *p* = 0.09) (Fig. 1 *left*).

**Fig. 1:**
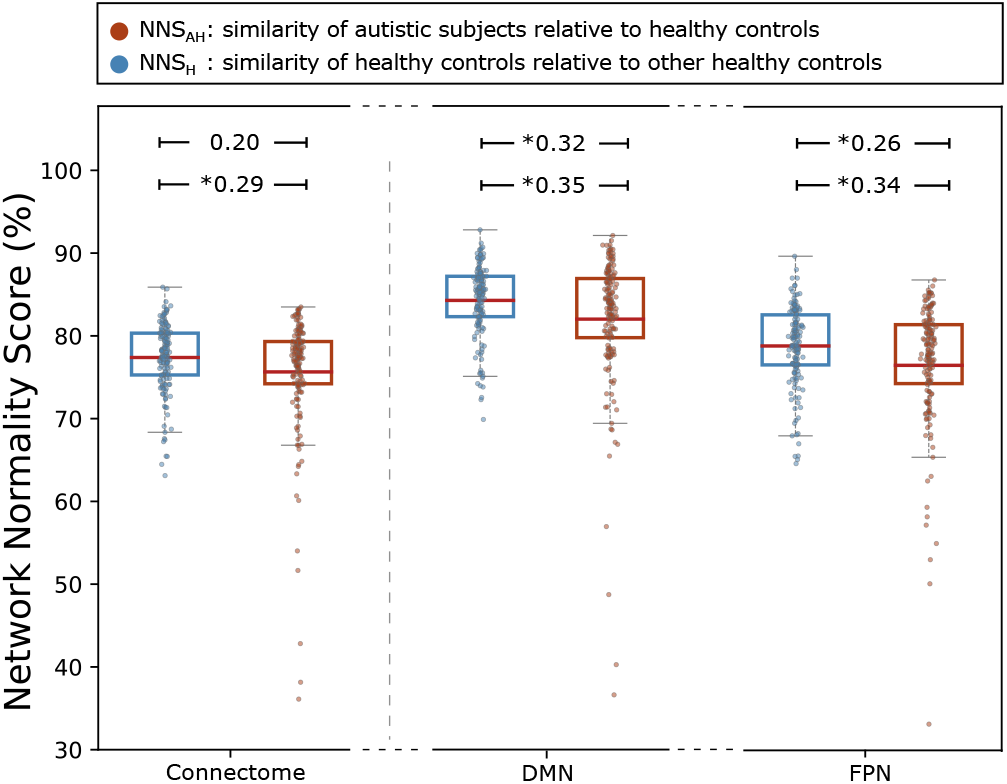
roup differences between NNS_H_ and NNS_AH_ at the connectome level (*Left*), DMN (*Middle*), and FPN (*Right*) levels. Effect sizes without and with outliers are shown on the first and second lines, respectively. Significant effect sizes are marked with a *. (*p* values are FDR corrected)

At the system level, we noted that the differences in structural connectivity between ASDs and HCs became particularly pronounced at the DMN (Fig. 1 *middle*) and FPN (Fig. 1 *right*). With outliers, we observed that NNS_AH_ was significantly lower than NNS_H_ in the DMN (*d* = 0.35, *p* = 0.004) and FPN (*d* = 0.34, *p* = 0.004). In the absence of outliers, we observed that NNS_AH_ was still significantly lower than NNS_H_ in the DMN (*d* = 0.32, *p* = 0.016) and FPN (*d* = 0.26, *p* = 0.038). These sub-network findings highlight a key topological difference in the structural connectivity of ASD subjects relative to the sample of healthy controls.

We then evaluated subgroups of Autism cohort by clustering ASDs into 2 clusters where three NNS_AH_ scores of individuals, namely connectome level and DMN and FPN system level scores, were utilized. (Fig. 2). We then evaluated relationship between NNS scores at connectome and system level for the clusters using the linear model (1).

**Fig. 2:**
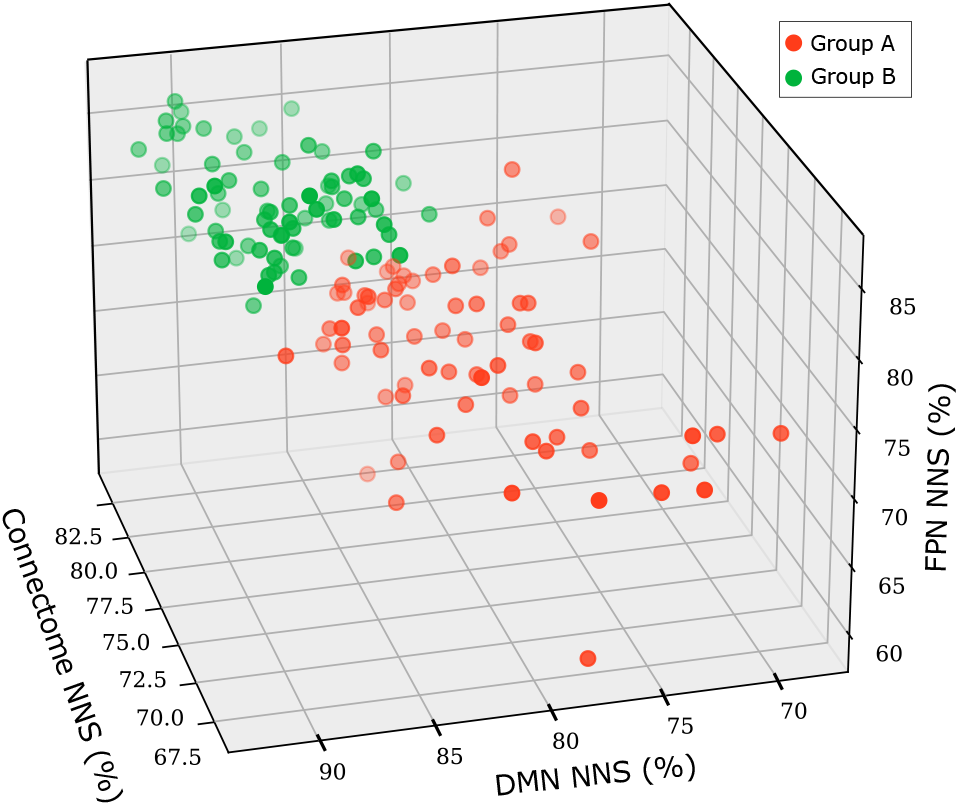
Sub-grouping of neuroimaging scores: Agglomerative clustering using NNSAH of connectome, DMN, and FPN levels to define two neuroimaging groups.

At the connectome level, we observed that NNS_AH_ and ADOS scores were significantly and negatively associated in group *A* (*r* = − 0.26, *p* = 0.036) (Fig. 3 *left*). At system level, DMN demonstrated a small negative association between NNS_AH_ and ADOS scores in group *B* (*r* = −0.38, *p* = 0.077) (Fig. 3 *right*), while FPN demonstrated strong negative associations between NNS_AH_ and ADOS scores in both groups: *A* (*r* = −0.38, *p* = 0.012) and *B* (*r* = −0.58, *p* = 0.030) (Fig. 4). A complete summary of results from the LM using these two clustered sub-groups is provided in Table 1.

**Table 1:** Results of fitting a linear model to evaluate the relationship between ADOS, NNS, and age. Estimated values by the linear model (which can be interpreted as correlations as terms were scaled) and p values are reported in each cell, respectively.

| | | NNS | Age | Adj. $R^2$ |
| --- | --- | --- | --- | --- |
| Connectome Level | Group A | **-0.26**<br>(0.036) | 0.18<br>(0.081) | 0.070 |
|  | Group B | -0.25<br>(0.290) | 0.29<br>(0.014) | 0.088 |
| Default Mode Network | Group A | -0.13<br>(0.369) | 0.18<br>(0.091) | 0.021 |
|  | Group B | -0.38<br>(0.077) | 0.30<br>(0.010) | 0.113 |
| Frontoparietal Network | Group A | **-0.38**<br>(0.012) | 0.17<br>(0.110) | 0.095 |
|  | Group B | **-0.58**<br>(0.030) | 0.31<br>(0.006) | 0.155 |

**Fig. 3:**
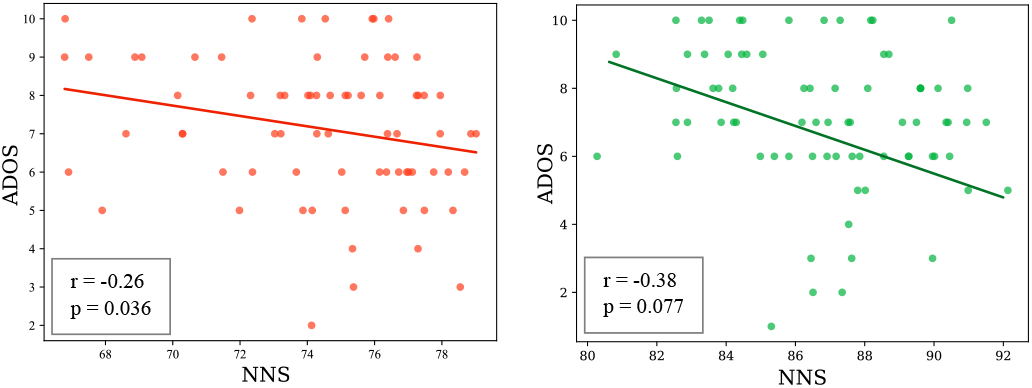
Relationship between NNS and autism severity. Using a LM, we observed a significant relationship between between NNSAH and ADOS for group *A* at connectome level (*Left*) and group *B* at DMN (*Right*).

**Fig. 4:**
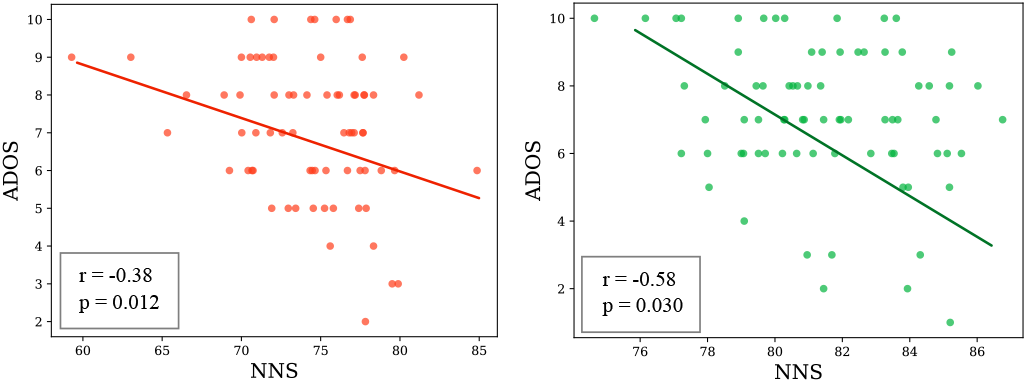
Relationship between NNS and autism severity. Using a LM, we observed a significant relationship between between NNSAH and ADOS for both group *A* (*Left*) and *B* (*Right*) at FPN.

## 4. DISCUSSION

In this study, we adopted a connectomic measure called network normality score (NNS) to Autism to demonstrate that structural network topology is altered in individuals with ASD compared to healthy controls. While we note that these differences are less prominent at connectome level, they are more pronounced in default mode and frontoparietal sub-networks. The DMN is a large-scale brain network that includes several high-level cognitive areas and is the portion of the brain functionally active even during resting state. fMRI studies have revealed that autistic brains exhibit aberrant patterns of functional connectivity in the DMN which have been linked with core ASD symptoms [9]. The frontoparietal network is responsible for coordinating behavior rapidly and accurately which is central to the completion of executive function tasks. Prior work has illustrated that regions of the frontal lobe and parietal cortex are involved in the mediation of impaired behaviors in autistic individuals [4]. While previous findings have shown these sub-network differences in functional connectivity, our findings indicate a structural basis for these altered network connectivity patterns in Autism.

We also described and quantified the association between brain network similarity of autistic individuals relative to healthy controls and their behavioral autism severity measured by ADOS. Establishing a relationship between the behavioral symptoms and neuroimaging measures is critical in understanding the etiology and neuropathology of the disorder. We showed significant associations in certain sub-groups between NNS_AH_ and ADOS at the whole connectome level, as well as at system level at DMN and FPN. This could indicate the efficacy of NNS in capturing the association between brain network dissimilarity and autism severity. While these findings are promising, we note that most of the associations discussed here being weak indicates a need for further investigation.

## 5. CONCLUSIONS

In this study, we have demonstrated the efficacy of our normative connectomic measure called NNS in capturing structural network alterations in ASD at connectome and system level. We showed that our findings of pronounced topological differences at the system level are consistent with established functional connectivity studies. Finally, we quantified the association between NNS and Autism severity to analyze its relationship with behavioral measures which provides insight into the underlying structural basis of the disorder.

## 6. COMPLIANCE WITH ETHICAL STANDARDS

This study was performed in line with the principles of the Declaration of Helsinki. Approval was granted by the Ethics Committee of Children’s Hospital Of Philadelphia.

## 7. ACKNOWLEDGEMENTS

No funding was received for conducting this study. The authors have no relevant financial or non-financial interests to disclose.

## REFERENCES

[1] Catherine Lord, Mayada Elsabbagh, Gillian Baird, and Jeremy Veenstra-Vanderweele, “Autism spectrum disorder,” The lancet, vol. 392, no. 10146, pp. 508–520, 2018.

[2] Christine Ecker and Declan Murphy, “Neuroimaging in autism—from basic science to translational research,” Nature Reviews Neurology, vol. 10, no. 2, pp. 82–91, 2014.

[3] Jinan Zeidan, Eric Fombonne, Julie Scorah, Alaa Ibrahim, Maureen S Durkin, Shekhar Saxena, Afiqah Yusuf, Andy Shih, and Mayada Elsabbagh, “Global prevalence of autism: a systematic review update,” Autism Research, vol. 15, no. 5, pp. 778–790, 2022.

[4] David G Amaral, Cynthia Mills Schumann, and Christine Wu Nordahl, “Neuroanatomy of autism,” Trends in neurosciences, vol. 31, no. 3, pp. 137–145, 2008.

[5] MR Herbert, D. Ziegler, CK Deutsch, L. O’brien, N Lange, A Bakardjiev, J Hodgson, KT Adrien, S Steele, N Makris, et al., “Dissociations of cerebral cortex, subcortical and cerebral white matter volumes in autistic boys,” Brain, vol. 126, no. 5, pp. 1182–1192, 2003.

[6] Mikail Rubinov and Olaf Sporns, “Complex network measures of brain connectivity: uses and interpretations,” Neuroimage, vol. 52, no. 3, pp. 1059–1069, 2010.

[7] Danielle S Bassett and Olaf Sporns, “Network neuroscience,” Nature neuroscience, vol. 20, no. 3, pp. 353– 364, 2017.

[8] Daniel A Abrams, Charles J Lynch, Katherine M Cheng, Jennifer Phillips, Kaustubh Supekar, Srikanth Ryali, Lucina Q Uddin, and Vinod Menon, “Underconnectivity between voice-selective cortex and reward circuitry in children with autism,” Proceedings of the National Academy of Sciences, vol. 110, no. 29, pp. 12060– 12065, 2013.

[9] Christopher S Monk, Scott J Peltier, Jillian Lee Wiggins, Shih-Jen Weng, Melisa Carrasco, Susan Risi, and Catherine Lord, “Abnormalities of intrinsic functional connectivity in autism spectrum disorders,” Neuroimage, vol. 47, no. 2, pp. 764–772, 2009.

[10] Naama Barnea-Goraly, Linda J Lotspeich, and Allan L Reiss, “Similar white matter aberrations in children with autism and their unaffected siblings: a diffusion tensor imaging study using tract-based spatial statistics,” Archives of general psychiatry, vol. 67, no. 10, pp. 1052–1060, 2010.

[11] Daniel H Geschwind and Pat Levitt, “Autism spectrum disorders: developmental disconnection syndromes,” Current opinion in neurobiology, vol. 17, no. 1, pp. 103– 111, 2007.

[12] Yusuf Osmanlioğlu, Drew Parker, Jacob A Alappatt, James J Gugger, Ramon R Diaz-Arrastia, John Whyte, Junghoon J Kim, and Ragini Verma, “Connectomic assessment of injury burden and longitudinal structural network alterations in moderate-to-severe traumatic brain injury,” Human brain mapping, 2022.

[13] Stacy Shumway, Cristan Farmer, Audrey Thurm, Lisa Joseph, David Black, and Christine Golden, “The ados calibrated severity score: relationship to phenotypic variables and stability over time,” Autism Research, vol. 5, no. 4, pp. 267–276, 2012.

[14] Yusuf Osmanlioğlu, Jacob A Alappatt, Drew Parker, and Ragini Verma, “Connectomic consistency: a systematic stability analysis of structural and functional connectivity,” Journal of neural engineering, vol. 17, no. 4, pp. 045004, 2020.

